# Combinatorial logic of Nav channels in nociceptor excitability: Different degrees of synergy define distinct neuronal groups

**DOI:** 10.64898/2026.04.04.716368

**Authors:** Dmytro Vasylyev, Sidharth Tyagi, Mohammad-Reza Ghovanloo, Peng Zhao, Stephen G. Waxman

## Abstract

Despite the genetic validation of voltage-gated sodium (Nav) channels as targets for pain therapy, single-channel subtype inhibition in clinical trials has yielded limited efficacy. Because Nav1.7 and Nav1.8 are thought to act in a coupled manner in ignition and subsequent regenerative, overshooting depolarization underlying the action potential of nociceptors, we utilized dynamic clamp, which permits precisely calibrated addition or subtraction of the current from any given channel, to systematically interrogate their combinatorial landscape. We demonstrate that nociceptor excitability is governed by highly non-linear biophysical rules. Partial, simultaneous attenuation of both conductances drives a supralinear collapse of action potential electrogenesis in sensory neurons. Unsupervised clustering reveals that this synergistic vulnerability is not a continuum, but varies in different neuronal subtypes: a population of sensory neurons experiences profound excitability suppression, whereas another population remains relatively resistant. These non-linear boundaries demonstrate that the efficacy of Nav current abrogation is fundamentally restricted by the underlying electrogenic architecture of the neuron, and provide a mechanistic blueprint for subtype-selective silencing.

## Introduction

Voltage-gated sodium (Nav) channels are among the most genetically and physiologically validated targets for pain therapy^1^. Within primary sensory neurons, Nav1.7 and Nav1.8 have emerged as especially important determinants of nociceptor excitability^2^. Nav1.7 amplifies small subthreshold depolarizations and helps set the gain for action potential initiation, while Nav1.8, by virtue of its depolarized availability and rapid recovery from inactivation, supports the regenerative action potential upstroke and repetitive firing of pain-signaling neurons^3,4^. The roles of Nav1.7 and Nav1.8 have been substantially genetically validated in a series of pain-related disorders^5–9^. Despite this strong target validation, clinical success with single-channel subtype inhibition has been incomplete. Selective Nav1.8 inhibition with VX-548 (Suzetrigine/Journavx) reduces acute postoperative pain in humans, but the magnitude of analgesia was partial^10,11^. Consistent with this, recent work in human dorsal root ganglion (DRG) neurons showed that even strong pharmacological inhibition of Nav1.8 reduces, but does not abolish repetitive firing, in part because excitability can still be supported by other sodium currents, especially Nav1.7^12–14^.

These observations suggest that the limited efficacy of monotherapy is not simply a pharmacokinetic, pharmacodynamic, or medicinal chemistry problem, but may reflect a deeper biophysical constraint imposed by the electrogenic architecture of nociceptors themselves. Prior dynamic clamp and modeling studies have shown that Nav1.7 and Nav1.8 do not function as isolated, interchangeable depolarizing currents. Rather, they play temporally and functionally distinct roles during action potential evolution, with Nav1.7 contributing strongly near resting voltages and Nav1.8 amplifying excitability over a broader depolarized range, particularly during repetitive firing^3,14,15^. In parallel, recent work demonstrated that subtraction of Nav1.8 conductance reveals a population of weak-responder DRG neurons in which even major loss of Nav1.8 does not produce strong suppression of excitability, indicating an intrinsic limit to the efficacy of Nav1.8-directed monotherapy^14^. If Nav1.7 and Nav1.8 operate as a coupled conductance system, however, it might be expected that the effect of perturbing both channels together cannot be inferred from single-channel perturbations alone. Under that framework, nociceptor excitability might obey nonlinear combinatorial rules rather than additive pharmacological logic.

This possibility is especially important in light of the marked heterogeneity of sensory neurons. Transcriptomic and cross-species atlas studies have revealed multiple molecularly distinct classes of DRG and trigeminal neurons, with differing complements of ion channels, neuropeptides, and sensory specializations^16–18^. Such diversity likely contributes to the fact that pain is not experienced as a unitary modality, but instead includes distinct percepts such as punctate mechanical pain, heat pain, spontaneous burning pain, and allodynia^19,20^. Although the neuronal substrates of these pain qualities are not yet fully resolved, the existence of multiple sensory neuron classes raises the possibility that different pain phenotypes recruit different electrophysiological clusters, each governed by different conductance rules. In that sense, transcriptomics and proteomics define what a sensory neuron is composed of, whereas systematic electrophysiological perturbation can begin to define how that neuron behaves as a dynamical excitable system. The present study is an initial step toward a functional electrophysiological-omics, or electromics, of nociceptor excitability.

To directly interrogate this problem, we used precisely calibrated dynamic clamp-mediated virtual subtraction of the Nav1.7 and Nav1.8 currents to construct a systematic 16-node perturbation matrix spanning discrete combinations of Nav1.7 and Nav1.8 attenuation in native DRG neurons. This approach allowed us to mimic idealized partial combinatorial block while preserving the endogenous membrane environment, and thus to ask whether paired reduction of the principal nociceptor sodium dyad produces merely additive effects or instead unmasks nonlinear vulnerability of the membrane. Dynamic clamp is uniquely suited to this question because it enables precise, real-time manipulation of specific conductances in intact neurons, allowing the combinatorial interaction landscape of excitability to be experimentally mapped rather than inferred^21–23^.

Here we show that partial simultaneous attenuation of Nav1.7 and Nav1.8 produces a supralinear collapse of action potential electrogenesis in a subset of sensory neurons, revealing that excitability is governed by nonlinear combinatorial logic. Importantly, unsupervised clustering analysis further demonstrates that this synergy is not universal and is not distributed as a continuum, but instead segregates neurons into distinct functional response classes. Our observation of populations of cells that are profoundly vulnerable and others that remain comparatively resistant even under strong dual subtraction identify a previously unrecognized multiplicity of excitability regimes within nociceptors and suggest that the efficacy of sodium channel targeting is fundamentally constrained by cluster-specific electrogenic architecture. More broadly, they imply that optimal analgesic strategies may require not only combined Nav1.7-Nav1.8 channel targeting, but also an understanding of which functional neuronal clusters are engaged in different pain states, and which additional conductances must be modulated to silence resistant populations.

## Results

### Systematic perturbation mapping reveals supralinear synergy between Nav1.7 and Nav1.8 in sensory neurons

To determine whether the functional interaction between Nav1.7 and Nav1.8 in DRG neurons is additive or non-linear (supra- or sub-additive), we used dynamic clamp to systematically subtract defined fractions of each conductance while recording excitability of that cell in current-clamp mode. This approach enabled us to move beyond single-channel perturbation and experimentally map the combinatorial conductance landscape governing action potential initiation. In each neuron, multiple recordings in current-clamp mode were taken, where cells were subjected to a 16-node perturbation matrix in which the two conductances were virtually subtracted in discrete combinations spanning 0, 25, 50, and 100% of each isoform (**Figure 1A**). Rheobase was then measured at each node as a direct readout of the current required to trigger an action potential. The experimental pipeline and baseline biophysical properties of the recorded neurons are summarized in **Supplementary Figure 1**.

**Figure 1.**
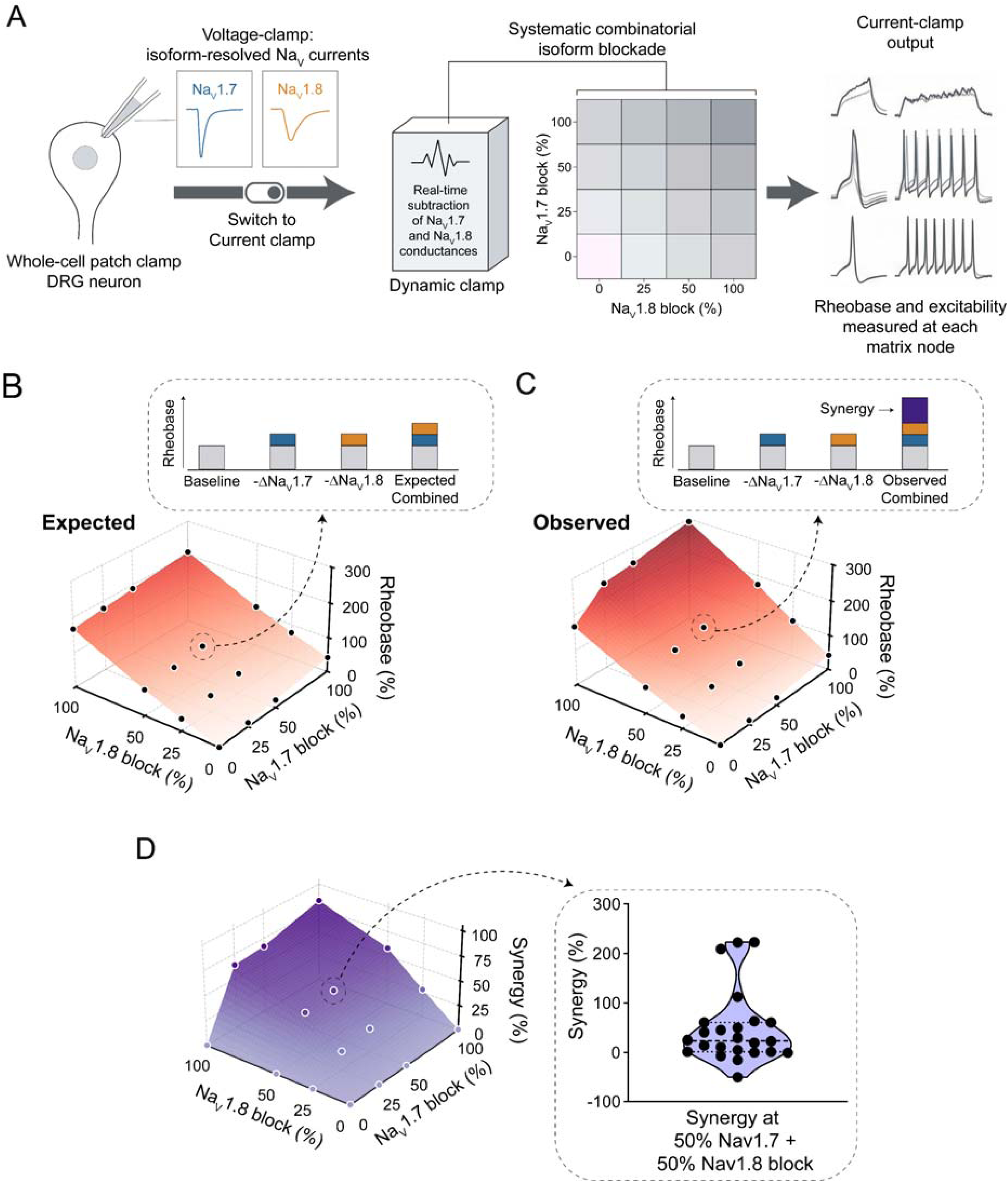
Systematic mapping of Nav1.7 and Nav1.8 combinatorial interactions via dynamic clamp. **(A)** Experimental workflow for dynamic clamp investigation of combinatorial isoform blockade in DRG neurons. Following voltage-clamp characterization of Nav1.7 and Nav1.8 currents, cells are switched to current-clamp mode and successive subtraction of specific conductances simulates systematic combinatorial blockade across a 16-node matrix **(B)** Representative 3D surface plot depicting the "Expected" rheobase at each matrix node, calculated by the linear summation of independent monotherapy penalties. Inset: Stacked bar chart illustrating the additive prediction for the 50/50 combination node. **(C)** Representative 3D surface plot of actual dynamic clamp recordings from the same neuron. The "Observed" rheobase significantly exceeds the additive prediction at combinatorial nodes, indicated by the purple "Synergy" component in the inset bar chart. **(D)** Violin plot showing the distribution of synergistic deviation (%) at the 50% + 50% Nav1.7+Nav1.8 node across all recorded neurons (n=24).

We first asked whether the effect of combined Nav1.7 and Nav1.8 attenuation could be predicted by simple linear summation of the corresponding monotherapy penalties. For each neuron, we therefore generated an additive expectation surface in which the rheobase shift at each combinatorial node was calculated from the independent effects of subtracting each isoform alone (**Figure 1B**). Under this framework, dual attenuation should increase rheobase in a manner that reflects only the arithmetic sum of the two single-channel perturbations. However, comparison of these expected surfaces with the actual dynamic clamp recordings revealed a markedly different behavior. In many neurons, the observed rheobase at combinatorial nodes substantially exceeded the additive prediction, producing a prominent upward deviation in the response surface that was absent from the corresponding monotherapy axes (**Figure 1C**). Subtraction of either Nav1.7 or Nav1.8 alone produced the expected increase in rheobase, but partial subtraction of both channels together often drove a far larger suppression of excitability than would be anticipated from either perturbation in isolation (**Figure 1D**). In these neurons, the membrane did not behave as though Nav1.7 and Nav1.8 were independent depolarizing reserves. Instead, the effect of attenuating one conductance depended strongly on the remaining availability of the other, indicating that the two channels operate as an electrogenic dyad functionally coupled in a nonlinear manner.

To quantify this interaction across the population, we calculated the percent synergistic deviation at each matrix node, defined as the excess rheobase shift above the additive expectation. We focused on the 50/50% subtraction node because it provided a balanced and physiologically informative measure of partial dual attenuation. At this node, the distribution of synergistic deviation across neurons was shifted above zero, demonstrating that the population response was dominated by supralinear rather than additive behavior (**Figure 1D**). Notably, the magnitude of this deviation varied substantially between cells, suggesting that although synergy is a general feature of the sensory neuronal Nav1.7-Nav1.8 system, its strength is not uniform across the population.

These results demonstrate that the DRG membrane obeys non-linear combinatorial rules of excitability. The consequences of jointly attenuating Nav1.7 and Nav1.8 cannot be inferred from single-channel perturbations alone. Instead, partial reduction of both conductances produces a supralinear collapse of action potential electrogenesis in a substantial fraction of sensory neurons. This observation establishes the existence of a functional interaction landscape within the nociceptor sodium dyad and raises the possibility that different neurons occupy distinct excitability regimes within that landscape.

### Unsupervised clustering identifies distinct functional excitability clusters with differential vulnerability to Nav1.7-Nav1.8 perturbation

The remarkable cell-to-cell variability in synergistic deviation observed in **Figure 1** suggested that the combinatorial logic of the Nav1.7-Nav1.8 dyad is not uniform across the DRG neuronal population. We therefore asked whether neurons could be partitioned into distinct functional classes on the basis of their full matrix responses rather than by any single biophysical parameter alone. To address this, we performed unsupervised spectral clustering analysis using the 16-node rheobase response vector from each neuron as the input feature space. This analysis resolved three discrete response classes, which segregated in principal component space into non-overlapping functional clusters, here termed C0, C1, and C2 (**Figure 2A**).

**Figure 2.**
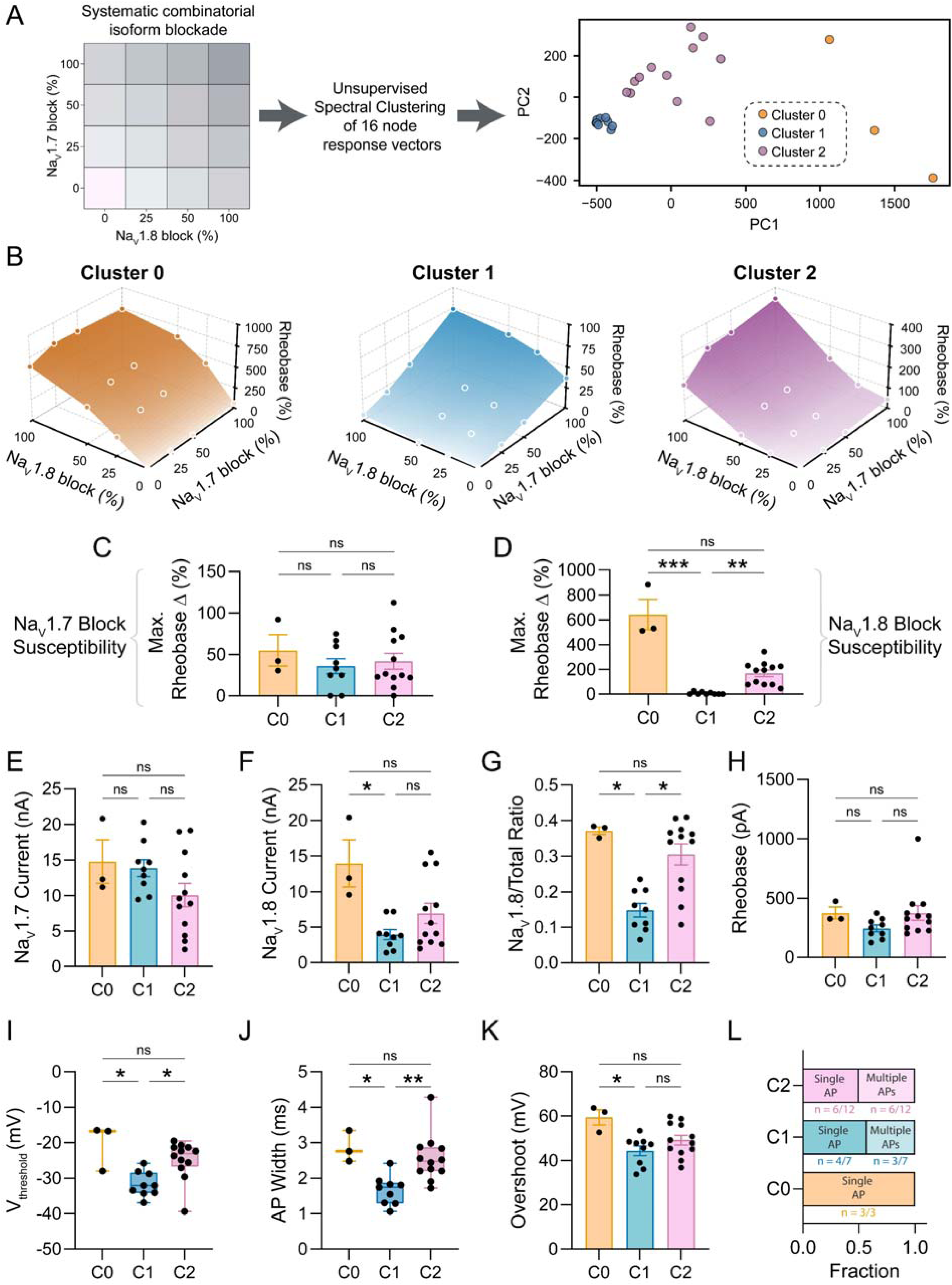
Unsupervised clustering reveals subtype-specific isoform blockade phenotypes. **(A)** Unsupervised spectral clustering of the 16-node response vectors and mapping to PCA space reveal three distinct functional clusters (C0, C1, C2). **(B)** Mean 3D rheobase surfaces for each cluster illustrate divergent sensitivities to the NaV dyad. Note the distinct z-axis scales, highlighting the high resistance of Cluster 1 compared to the extreme sensitivity of Clusters 0 and 2 to Nav1.8 blockade. **(C)** Maximum rheobase delta in response to Nav1.7 subtraction, stratified by cluster **(D)** Maximum rheobase delta in response to Nav1.8 subtraction, stratified by cluster **(E)** Comparison of isolated NaV1.7 currents between the clusters **(F)** Comparison of isolated NaV1.8 currents between the clusters **(G)** Comparison of NaV1.8/Total sodium currents between the clusters **(H)** Comparison of Baseline Rheobase between the clusters **(I)** Comparison of Vthreshold between the clusters **(J)** Comparison of action potential halfwidth between the clusters **(K)** Comparison of Action potential overshoot between the clusters **(L)** Proportion of neurons in each cluster exhibiting single vs. multiple action potential firing in response to current injection.

### Clusters diverge not in Nav1.7 dependence, but in their integrated response to Nav1.8 loss and dual perturbation

The average rheobase surfaces for these clusters revealed fundamentally different interaction architectures (**Figure 2B**; **Supplementary Figure 2**). Cluster 1 displayed a comparatively shallow surface, indicating broad resistance to perturbation of the Nav dyad. In contrast, Clusters 0 and 2 showed much steeper surfaces, consistent with a substantially greater rise in firing threshold across the matrix, and thus greater vulnerability to conductance subtraction. Importantly, these differences were not captured simply by comparing responses to one channel in isolation. Subtraction of Nav1.7 alone produced relatively similar rheobase shifts across clusters (**Figure 2C**), whereas subtraction of Nav1.8 exposed a much more pronounced divergence, with one population showing strong sensitivity and another exhibiting relative insensitivity (**Figure 2D**). Thus, the major distinction between clusters did not lie in whether Nav1.7 contributed to excitability, but in how the membrane integrated Nav1.8 loss and, more importantly, how it behaved when both conductances were perturbed together.

We next asked whether these functional clusters corresponded to differences in baseline sodium current properties. Comparison of isolated Nav1.7 currents showed no simple monotonic relationship that could fully explain the clustering pattern (**Figure 2E**), whereas isolated Nav1.8 current and the Nav1.8-to-total sodium current ratio differed more clearly between groups (**Figure 2F-G**). These relationships are consistent with the idea that Nav1.8-rich neurons are more vulnerable to perturbation of the dyad. However, Nav1.8-related measures alone were insufficient to account for the full clustering structure (**Supplementary Figure 2**). This is important because it indicates that the functional response classes reflect integrated electrogenic architecture rather than a simple scalar continuum of Nav1.8 functional expression. In other words, the clusters are not merely “more Nav1.8” versus “less Nav1.8” cells. Instead, they represent distinct classes of excitability behavior revealed only when this excitability is challenged across the full combinatorial matrix.

Baseline electrophysiological properties further supported this interpretation. Cluster membership was associated with differences in rheobase and voltage threshold (**Figure 2H-I**), as well as action potential width and overshoot (**Figure 2J-K**), indicating that the clusters occupy distinct baseline excitability states even before virtual channel subtraction. In particular, the broader-spiking populations were more vulnerable to dual-channel perturbation, whereas narrow-spiking neurons were comparatively resistant. These changes were also reflected in the response vectors of action potential morphology parameters (**Supplementary Figures 4-5**). By contrast, the proportion of neurons exhibiting single versus multiple action potentials did not differ significantly across groups (**Figure 2L**), suggesting that firing pattern alone was insufficient to distinguish the identified clusters. Together, these observations suggest that synergy emerges from the integrated electrophysiological phenotype of the cell, rather than from any single conductance measurement in isolation. Thus, waveform shape and firing behavior report the underlying electrogenic context in which the Nav1.7-Nav1.8 interaction is embedded.

Taken together, these data show that supralinear vulnerability to Nav1.7-Nav1.8 attenuation is highly subtype-contingent. The sensory neuronal population contains at least three functional excitability phenotypes, including clusters that are highly susceptible to combined perturbation and another that remains intrinsically resistant. This cluster-specific organization provides a mechanistic basis for why single-target and even dual-target sodium channel strategies may not produce uniform silencing across the entire population of nociceptors. It also suggests that sensory neurons can be classified not only by their molecular composition, but by the nonlinear conductance rules that govern their excitability.

### Action potential follows distinct cluster-specific trajectories during combined Nav1.7-Nav1.8 attenuation

We next asked whether the supralinear interaction identified above was organized similarly across all functional clusters, or whether each cluster occupied a distinct region of the combinatorial excitability landscape. Cluster-averaged synergy surfaces showed clear divergence between groups (**Figure 3A**). Clusters 0 and 2 displayed prominent synergy peaks within the interior of the matrix, indicating that partial simultaneous subtraction of Nav1.7 and Nav1.8 produced a disproportionately large suppression of excitability. In contrast, Cluster 1 showed a relatively flat surface, consistent with a resistant phenotype. We previously showed that rheobase penalties resulting from Nav1.8 abrogation in nociceptive neurons are correlated with the relative functional contributions of Nav1.7 and Nav1.8 in the perithreshold region^14^. In the present study, cluster 1 neurons exhibited hyperpolarized voltage thresholds and a low Nav1.8/Nav1.7 ratio in the perithreshold region, whereas voltage thresholds in cluster 0 and cluster 2 neurons were relatively depolarized, resulting in a high Nav1.8/Nav1.7 ratio in the perithreshold region (**Figure 3B-J**). Consistent with these observations, the rheobase penalty in cluster 1 neurons was small at the principal diagonal of the Nav1.7:Nav1.8 combinatorial matrix, whereas the corresponding penalty in cluster 0 and cluster 2 neurons was significantly larger (**Figure 3K-M**). Thus, the key insight from this analysis is that synergy is not merely present or absent across the population, but is organized in a cluster-specific manner that reflects distinct underlying electrogenic architectures.

**Figure 3.**
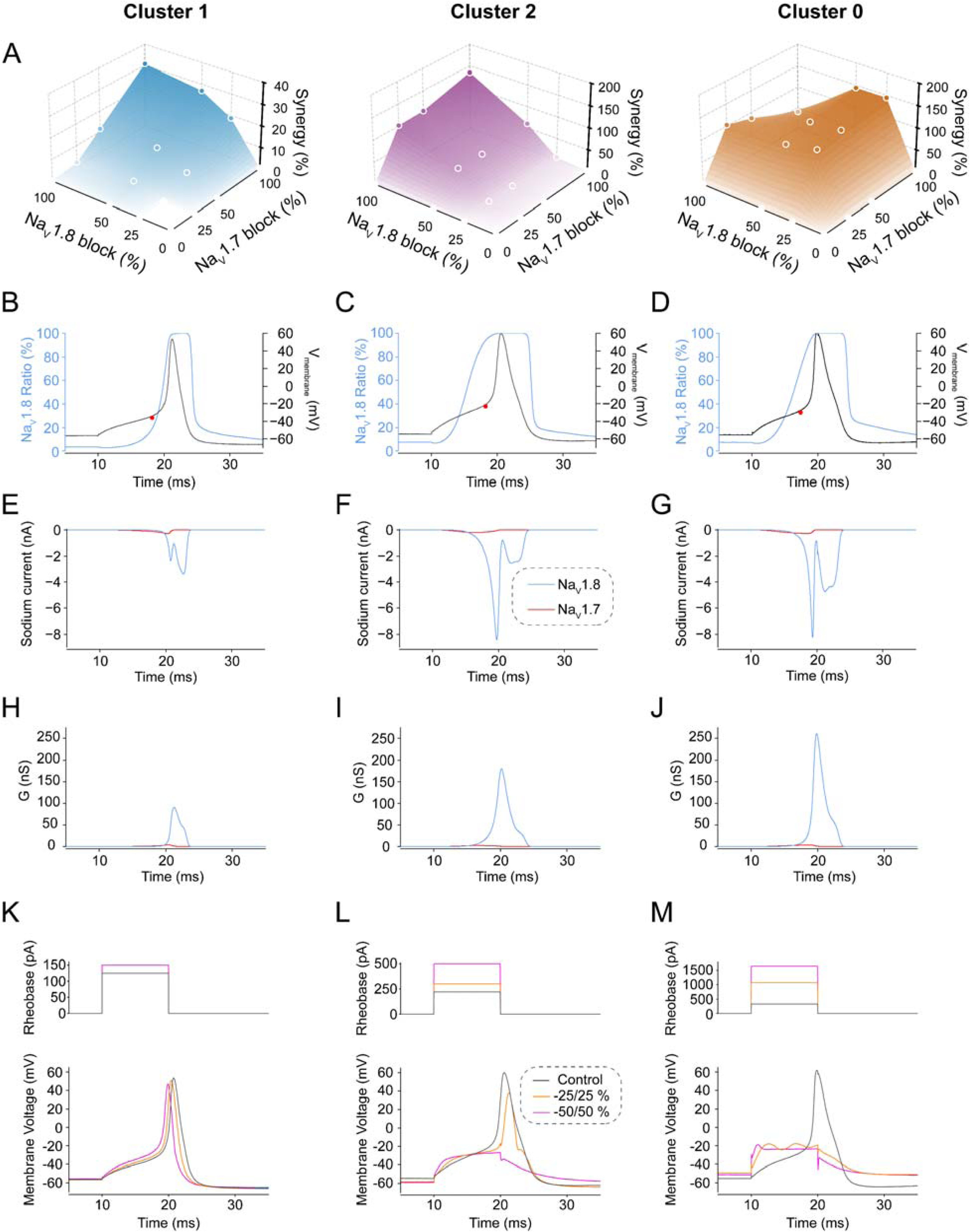
Action potential dynamics and synergistic differences between sensory neuronal clusters. **(A)** 3D surfaces mapping the percentage of synergy across the matrix for Clusters 1, 2, and 0. Clusters 2 and 0 exhibit a massive synergistic "peak" compared to the relatively flat profile of Cluster 1. (**B-D**) Current clamp recordings in small DRG neurons representing subpopulation of cluster 1 (**B**), cluster 2 (**C**) and cluster 0 (**D**) cells. APs were evoked by a 10-ms long current pulse at rheobase intensity (**K-M**, upper panel, black stimulus trace). The red dots indicate voltage thresholds of AP generation. Blue line (blue Y-axis) denotes relative contribution of Nav1.8 to the sum of Nav1.7 and Nav1.8 sodium currents calculated for the respective APs. Nav1.7 (red trace) and Nav1.8 (blue trace) sodium currents (**E-G**) and their respective conductances (**H-J**) were calculated using in-silico action potential clamp for the respective traces of membrane voltage shown in (**B-D**). Sodium currents were modeled based on Hodgkin-Huxley equations and calculated post hoc by APsim software using previously recorded APs, with the following Gmax values: cluster 1 neuron (**B**) Nav1.7 Gmax = 736 nS and Nav1.8 Gmax = 277 nS; cluster 2 neuron (**C**) Nav1.7 Gmax = 800 nS and Nav1.8 Gmax = 441 nS; cluster 0 neuron (**D**) Nav1.7 Gmax = 1034 nS and Nav1.8 Gmax = 642 nS. (**K-M**) Shown are current clamp recordings of action potentials (bottom panel) illustrating the rheobase penalty (rheobase stimulus is shown on top panel) at principal diagonal of the Nav1.7:Nav1.8 combinatorial matrix (control, black trace; 25%:25%, red trace; 50%:50%, blue trace) for the cluster 1 (**K**), cluster 2 (**L**) and cluster 0 (**M**) neuron (recordings are from the same neurons as reported in **B-D**).

### Non-linear interaction boundaries reveal dose-sparing efficacy but also an intrinsic limit of dual Nav1.7-Nav1.8 targeting

To assess the translational implications of the cluster-specific interaction landscape, we next asked how much Nav1.7 and Nav1.8 subtraction was required to produce a defined increase in firing threshold. We therefore constructed isobolograms for a 2-fold increase in rheobase, using this as a stringent benchmark for strong suppression of excitability. At the population level, the observed isobole bowed markedly inward relative to the additive expectation, indicating substantial dose-sparing synergy between Nav1.7 and Nav1.8 attenuation (**Figure 4A**). Thus, the same rheobase endpoint could often be reached with considerably less subtraction of each individual conductance when both were reduced together.

**Figure 4.**
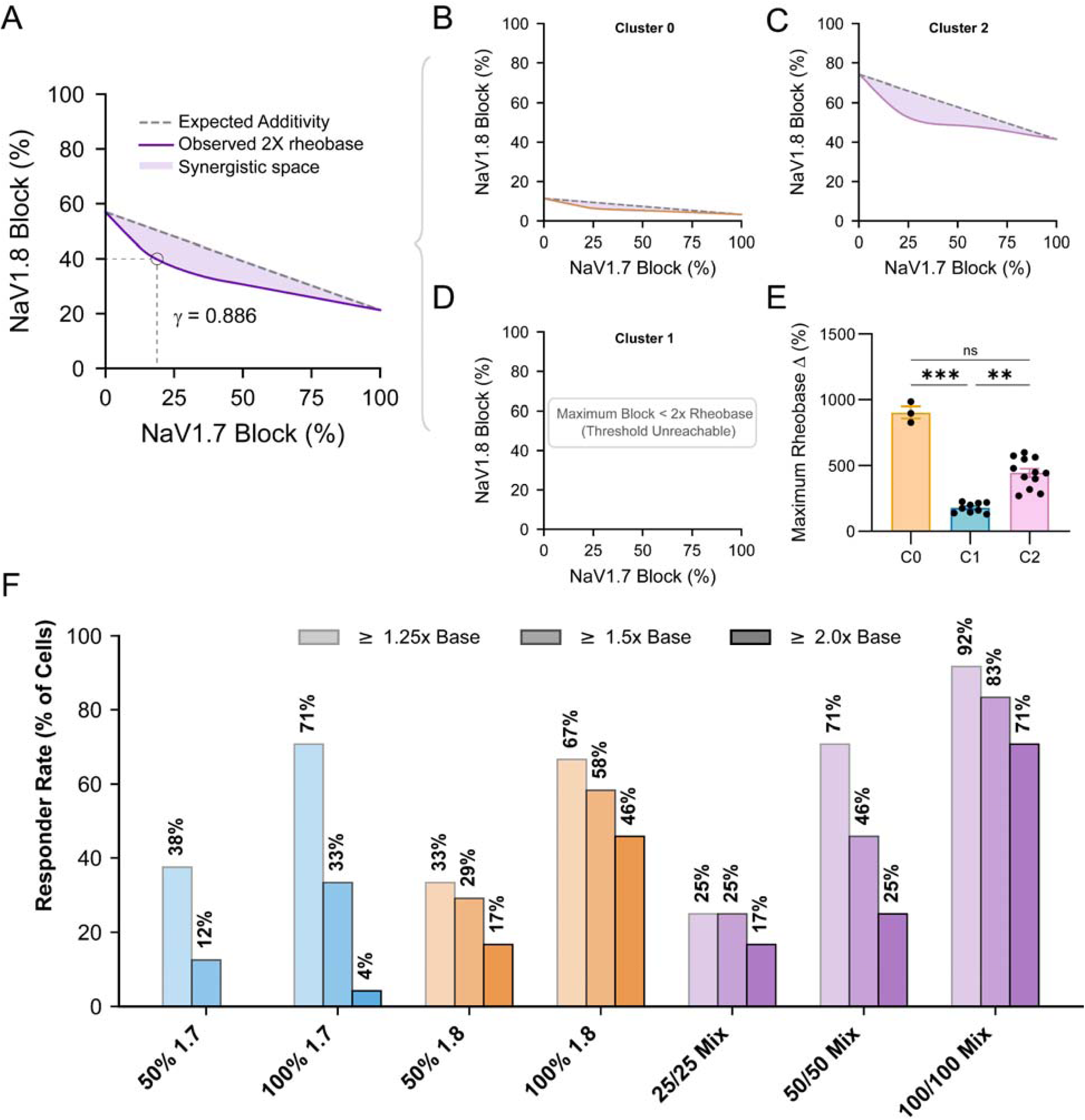
The potential of Nav1.7/Nav1.8 synergy for dose-sparing analgesia. **(A)** Population-level isobologram showing the percent reduction in Nav1.7 and Nav1.8 required to achieve a 2x increase in rheobase. The deep inward bow of the purple "Observed" curve relative to the "Expected" dashed line highlights the significant synergistic space. (**B-D**) Side-by-side comparison of 2x rheobase isoboles for Cluster 0 (**B**), Cluster 2 (**C**), and Cluster 1 (**D**). While Cluster 2 neurons show massive dose-sparing, the 2x threshold remains unreachable for Cluster 1 neurons even at 100% Nav1.7 or Nav1.8 blockade. (**E**) Comparison of the maximum achievable rheobase shift for each neuron, stratified by cluster. (**F**) Bar chart illustrating the percentage of cells that experienced a rheobase increase (at 1.25x, 1.5x, and 2.0x base thresholds) across isoform specific subtraction ratios.

When stratified by cluster, however, this synergistic space differed sharply across neuronal classes. Clusters 0 and 2 exhibited pronounced inward-bowed isoboles, consistent with strong combinatorial vulnerability and efficient dose-sparing (**Figure 4B-C**). In contrast, Cluster 1 remained comparatively resistant, and in many cells the 2-fold rheobase threshold was not reached even at maximal subtraction of either Nav1.7 or Nav1.8 alone, or under large, combined reductions of both conductances (**Figure 4D**). Thus, while combined targeting greatly increased efficacy in vulnerable clusters, it did not produce universal silencing across the population.

This cluster-dependent limit was further evident when the maximal achievable rheobase shift was compared across neurons (**Figure 4E**). Vulnerable clusters showed substantially larger dynamic ranges of suppression, whereas resistant neurons remained bounded by a lower ceiling of excitability reduction. Consistent with this, the fraction of cells achieving progressively stronger rheobase shifts varied across subtraction ratios and depended strongly on cluster identity (**Figure 4F**).

Together, these results show that dual Nav1.7-Nav1.8 targeting creates a substantial dose-sparing therapeutic space, but that this space is constrained by the underlying electrogenic architecture of the neuron. In some sensory neuronal clusters, partial combinatorial attenuation is sufficient to strongly suppress firing. In others, even major dual subtraction fails to drive equivalent silencing, revealing an intrinsic biophysical limit to this specific channel-targeting strategy.

## Discussion

Our results show that peripheral sensory neuron excitability is governed by non-linear combinatorial rules within the Nav1.7-Nav1.8 system. Although both channels are established contributors to nociceptor firing, clinical studies to date have treated them as partly independent pharmacological targets. Our perturbation mapping argues against that view. Partial simultaneous attenuation of Nav1.7 and Nav1.8 produced a supralinear increase in rheobase in a substantial fraction of neurons, indicating that the effect of reducing one conductance depends strongly on the remaining availability of the other. Thus, the membrane does not behave as a linear sum of depolarizing reserves, but as an integrated electrogenic system.

This work extends prior studies that defined complementary roles of Nav1.7 and Nav1.8 in DRG neurons. Nav1.7 contributes prominently near resting and perithreshold voltages, whereas Nav1.8, while contributing at perithreshold voltages in a population of neurons^14^, generally supports the suprathreshold regenerative and repetitive phases of firing^3,4^. More recent work showed that Nav1.8-directed perturbation alone encounters an intrinsic ceiling because some neurons remain weak responders^12–14^. The advance here is that mapping the dyad itself reveals a spectrum of non-linear interaction regimes that cannot be inferred from single-channel perturbations alone.

A central finding is that DRG neurons can be divided into discrete clusters on the basis of their Nav1.7-Nav1.8 interactions, and this synergy is cluster-dependent. Some neurons were profoundly vulnerable to combined Nav1.7-Nav1.8 subtraction, whereas another population remained comparatively resistant. This indicates that efficacy is constrained not simply by the abundance of one isoform, but by the broader electrogenic architecture of the neuron. In that sense, the identified clusters are functional entities defined by distinct rules of excitability.

This cluster-based view may also help frame pain heterogeneity. Transcriptomic and proteomic studies have established that peripheral sensory ganglia contain multiple molecularly distinct neuronal classes^16–18,24,25^. Microneurography, which permits recording from peripheral sensory axons in human subjects, does not provide direct information about the membrane properties of recorded neurons, but has identified multiple distinct classes of C fibers based on their functional response properties and activity-dependent slowing^26,27^. Our data add an electrogenic layer of DRG neuron differentiation by showing that neurons also differ in the combinatorial conductance logic that governs their excitability. We do not assign specific pain qualities to specific clusters here. However, the results raise the possibility that different pain phenotypes, such as allodynia, punctate pain, or heat pain, may recruit neuronal populations with different electrophysiological vulnerabilities. In this sense, systematic perturbation mapping can be viewed as an initial step toward a functional omics of excitability.

The translational implications of this work are twofold. First, combined attenuation of Nav1.7 and Nav1.8 creates a substantial dose-sparing space, suggesting that dual targeting of both channels may suppress excitability more effectively than selective inhibition of either isoform alone. This suggests a combinatorial approach to the incomplete efficacy of selective Nav1.8 inhibition that has been seen in both clinical studies and human DRG recordings^11,12^. Achieving this therapeutic dose-sparing space may not only involve combinations of traditional small molecules, but could also be realized through emerging modalities, such as gene therapy or targeted protein degradation^28^. However, even dual targeting of Nav1.7 and Nav1.8 was not sufficient to silence all neurons. A resistant cluster remained bounded by a limited maximal rheobase shift, revealing an intrinsic biophysical limit to this strategy. Thus, combined Nav1.7-Nav1.8 targeting should be expected to outperform monotherapy in some neuronal populations, but not to produce universal suppression.

These findings may also have implications for current therapeutic strategy. Recent analgesic development has increasingly emphasized precision targeting of one, or at most two, sodium channel isoforms. Our results support the logic of such approaches but also expose their likely limits. If distinct sensory neuron clusters are maintained by partially overlapping but non-identical conductance architectures, then even highly selective dual targeting may leave resistant populations insufficiently suppressed. In that context, safe and tolerable compounds that dampen excitability through actions at multiple ion channels and receptors may, in some settings, offer a more effective route toward broader pain control^29^. Nonpsychotomimetic cannabinoids are one example of such a class, since they can modulate excitability through Nav channels as well as additional molecular targets^30–34^. As a second example, lacosamide, which is functionally selective for inactivated channels but not subtype-specific, has been shown to ameliorate pain in peripheral neuropathy^35^. This does not argue against precision target-specific pharmacology, but suggests that for diverse pain states, especially those likely to recruit multiple sensory neuronal clusters, rationally designed or carefully selected polypharmacology may prove advantageous.

One likely explanation for the resistant phenotype is that action potential generation in these neurons is buffered by additional conductances beyond the Nav1.7-Nav1.8 dyad, including other inward sodium currents, high voltage-activated calcium current^36^ and/or differences in stabilizing outward currents^37,38^. In that framework, broader suppression may require targeting additional conductances^35^. Identification of those mechanisms will require further study.

This study has limitations. The identified clusters are electrogenic; further work will be needed for mapping onto transcriptomic classes or specific sensory modalities. The experiments were performed under controlled *in vitro* conditions, and the matrix necessarily interrogates only two conductances within a broader excitability network. Even so, the present results are important in directly revealing non-linear rules of excitability that are inaccessible to single-channel perturbation alone, in demonstrating the presence of electrogenically distinct clusters of neurons, and in indicating that even full abrogation of Nav1.7 and Nav1.8 will not mute the hyperactivity of the full population of DRG neurons.

A logical next step will be to extend this framework beyond Nav1.7 and Nav1.8 by mapping other ion channels across the defined functional clusters. Doing so should help build a richer atlas of sensory neuron functional -omics (electromics), linking cluster-specific excitability rules to the broader conductance ensembles that shape firing behavior. Integration of this type of electrophysiological perturbation mapping with transcriptomic and molecular profiling in physiologically normal and pathological states may ultimately provide a more complete classification of sensory neurons, not only by what molecules they express, but by how their excitability is organized and how it can be therapeutically modulated^39^.

In summary, our results show that nociceptor excitability is governed by cluster-specific non-linear combinatorial logic within the Nav1.7-Nav1.8 dyad. Partial dual attenuation can produce profound supralinear suppression in some neurons, yet even strong abrogation of both currents remains insufficient in others. These findings provide a functional framework for understanding why sodium channel-targeting strategies succeed in some neuronal populations and fail in others, and suggest that future analgesic strategies may need to be matched to the functional excitability classes engaged in different pain states.

## Acknowledgments

The Center for Neuroscience and Regeneration Research is a Collaboration of the Paralyzed Veterans of America with Yale University. We thank Joshua Huttler for helpful discussions.

## Author contributions

D.V., S.T., M.-R.G., and S.G.W. contributed to design; D.V. and P.Z. performed research; D.V., S.T., and M.-R.G., interpretation; D.V., patch-clamp; D.V., S.T., analysis; S.T. performed computational analyses; S.G.W. supervised the study; M.-R.G. and S.T. prepared the first draft; and all authors edited and approved the final manuscript.

## Methods

### DRG neuron preparation

Time-pregnant dams were purchased from Inotiv/Envigo and arrived in-house at E17-E18. Pups were maintained with the respective dam, who had free access to food and water. DRGs from male postnatal day 4 – 5 (P4 – P5) Sprague–Dawley rats were harvested and dissociated as reported previously^40^. Briefly, DRGs were harvested and placed in ice-cold complete saline solution (CSS) [in mM: 137 NaCl, 5.3 KCl, 1 MgCl2, 25 sorbitol, 3 CaCl2, and 10 N-2-hydroxyethylpiperazine-N′-2-ethanesulfonic acid (HEPES), adjusted to pH 7.2 with NaOH]. DRGs were then incubated at 37°C for 20-min in CSS containing 1.5 mg/ml Collagenase A (Roche) and 0.6 mM EDTA, followed by 20-min incubation at 37°C in CSS containing 1.5 mg/ml Collagenase D (Roche), 0.6 mM EDTA, and 30 U/mL papain (Worthington Biochemical); the DRGs were then triturated in DRG culture media, supplemented with 1.5 mg/ml low-endotoxin BSA (Sigma) and 1.5 mg/ml trypsin inhibitor (Sigma). DRG culture media was DMEM/F12 (Invitrogen) supplemented with 10% fetal bovine serum (Hyclone), 100 U/ml penicillin, 0.1 mg/ml streptomycin (Invitrogen), and 2 mM L-glutamine (Invitrogen). After trituration, 100 µl of cell suspension was seeded directly onto each poly-D-lysine/laminin-coated coverslip (Corning, Discovery Labware) and incubated at 37°C in a 95% air/5% CO2 (vol/vol) incubator. After allowing 45 min for neurons to attach to the coverslips, DRG media supplemented with 50 ng/ml NGF and 50 ng/ml GDNF was added into each well to a final volume of 1.0 ml and the neurons were maintained at 37°C in a 95% air/5% CO2 (vol/vol) incubator until used for electrophysiological experiments. Half of the media supplemented with 50 ng/ml NGF and 50 ng/ml GDNF was changed every other day.

### Electrophysiology and dynamic clamp

Small-diameter DRG neurons (soma diameter 24-28 μm, 25.88 ± 1.1 µm, mean ± SD, n = 24) were studied in whole-cell configuration under dynamic clamp^3,22,41–46^. Membrane voltage and current recordings were acquired using a HEKA EPC10 USB amplifier and PatchMaster software interfaced with a Cybercyte CIM/V10 dynamic clamp system with 20–25 μs latency. Voltage and current traces were filtered at 3 kHz and digitized at 50 kHz. Recordings were performed at room temperature (21–23 °C). Patch pipettes had resistances of 1.5–3 MΩ when filled with intracellular solution containing (in mM): 150 KCl, 0.5 EGTA, 5 HEPES, 3 Mg-ATP, and 5.6 glucose, adjusted to pH 7.3 with KOH, 286-288 mOsm. Extracellular solution was HBSS (Invitrogen, catalog number 14025) (in mM): 1.3 CaCl_2_, 0.5 MgCl_2_, 0.4 MgSO_4_, 5.3 KCl, 0.4 KH_2_PO_4_, 4.2 NaHCO_3_, 138 NaCl, 0.3 Na_2_HPO_4_, 5.6 glucose, 284-286 mOsm. During current-clamp recordings, neurons were maintained at their native resting membrane potential without steady-state current injection. Stability of the recordings (rheobase, RMP, overshoot) was assessed at the end of each matrix column (every four recordings) by repeating the measurements under control conditions. Rheobase was measured using 10-ms depolarizing current injections, and voltage threshold was defined as the point at which the first derivative of the membrane potential began to rise sharply from baseline. Baseline electrophysiological properties (Supplementary Figure 1), including resting membrane potential, voltage threshold, action potential width, overshoot, undershoot, input resistance, and firing pattern, were also quantified.

For each neuron, whole-cell voltage clamp was first used to estimate endogenous Nav1.7 and Nav1.8 current amplitudes. Nav1.8 current was isolated using depolarizing voltage steps from a holding potential of −45 mV, at which most other sodium channel isoforms are inactivated while Nav1.8 remains available. Nav1.7 current was estimated from the tetrodotoxin-sensitive sodium current component based on prior evidence that Nav1.7 constitutes ∼70% of tetrodotoxin-sensitive current in small DRG neurons^45^. These experimentally measured sodium current amplitudes were then used to scale Hodgkin-Huxley-based Nav1.7 and Nav1.8 models within the dynamic clamp system by adjusting the maximal conductance (Gmax) for each channel model so that modeled currents matched the magnitude of the endogenous sodium currents recorded from that cell. Dynamic clamp current injected through the recording pipette therefore functionally mimicked the amplitude and timing of native sodium currents while preserving endogenous channel expression and activity.

The kinetic models utilized in this study were derived from the Hodgkin-Huxley framework^47^, following the equation dx/dt = α_x_(1-x) - β_x_x, where *x* represents the channel gating variable, and α and β are the forward and backward rate constants (ms⁻¹). Sodium current was expressed as I_Na_ = G_max_*m^3^*h*s*(V_m_-E_Na_), where G_max_ is the maximum conductance, V_m_ is membrane voltage, and E_Na_ (sodium reversal potential) was set at 65 mV. The activation variable is denoted as *m*, while *h* and *s* represent inactivation variables (*s* = 1 for Nav1.8 channels).

To simulate sodium currents, we applied in silico action potential clamp (AP clamp) protocols^36,45^, computing responses offline at 20 μs resolution to match PatchMaster digitization protocol. Calculations were performed using APsim, a custom program scripted in LabTalk within the Origin framework^45^. Hodgkin-Huxley variables, their products (m^3^hs), and corresponding ionic currents were generated computationally using APsim. In silico modeling of Nav1.7 and Nav1.8 currents was conducted by matching the model’s maximal conductance values to the respective native currents recorded from each cell. The model’s conductance was adjusted to ensure that the peak amplitude of simulated currents aligned with experimentally recorded values under identical voltage protocols. Values of Gmax for Nav1.7 and Nav1.8 are provided in the text as 100% conductance.

The Nav1.7 channel model was from Vasylyev et al. 2014^45^:

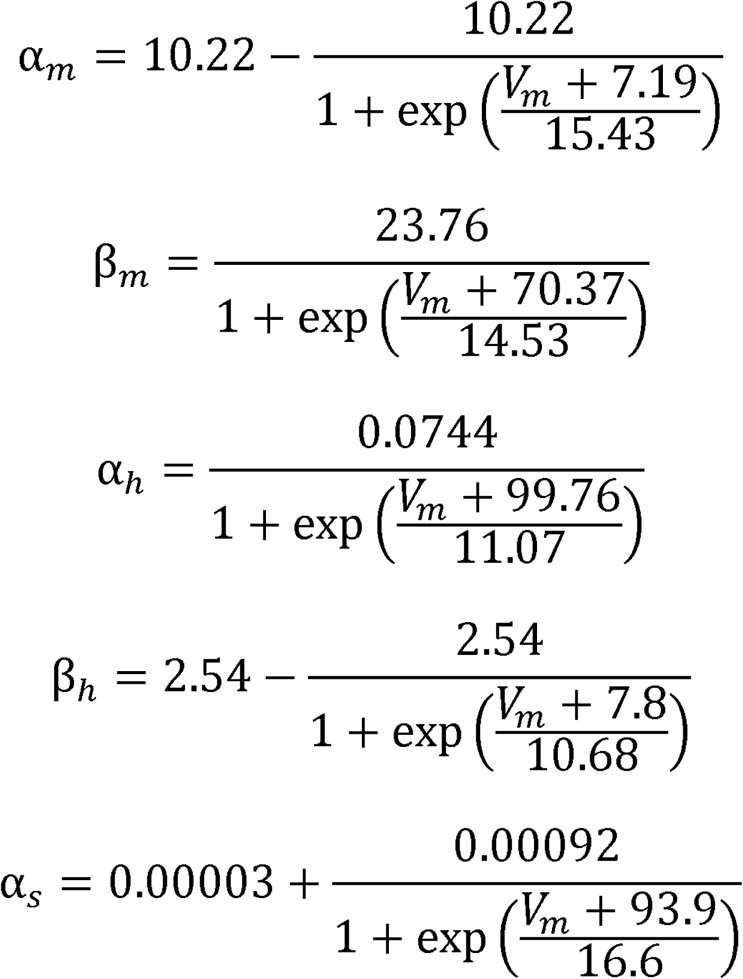

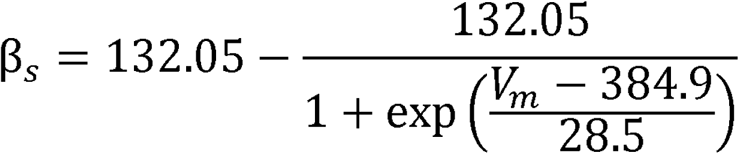

The Nav1.8 channel model was from Sheets et al. 2007^48^:

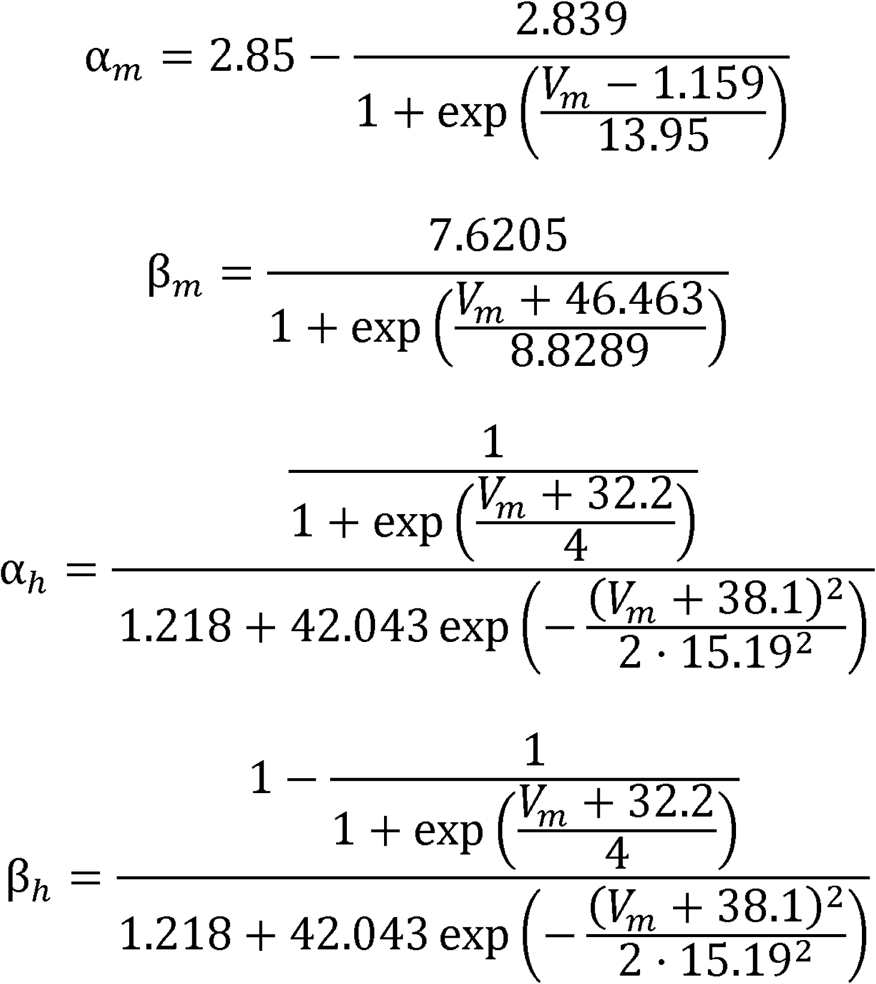

### Quantification of synergy

For each neuron, an additive expectation surface was calculated from the independent rheobase penalties produced by single-axis Nav1.7 or Nav1.8 subtraction. Synergy was defined as the excess observed rheobase shift above this additive prediction at each combinatorial node and was expressed as percent deviation from expected.

### Clustering and computational analysis

For clustering analyses, each neuron was represented by a 16-dimensional rheobase response vector corresponding to all combinations of Nav1.7 and Nav1.8 subtraction (0, 25, 50, and 100% for each conductance). Features were arranged in a fixed order with Nav1.8 subtraction varying along the outer dimension and Nav1.7 subtraction varying along the inner dimension, such that the vector corresponded to the full 4 × 4 subtraction matrix from (0% Nav1.7, 0% Nav1.8) through (100% Nav1.7, 100% Nav1.8). The rheobase feature used for clustering was the baseline-normalized rheobase value, which was anchored at 100 for the control condition.

Unsupervised clustering was performed using spectral clustering on the raw, unscaled feature matrix, with three clusters specified a priori based on qualitative inspection of the data and to balance separation of distinct response phenotypes against overfragmentation of the dataset. Spectral clustering used a nearest-neighbor affinity graph with 10 neighbors, selected because this value produced a fully connected graph and relatively stable cluster assignments across a range of neighbor values. Principal component analysis (PCA) was used to visualize clustering structure in two dimensions. Cluster-average heatmaps of rheobase responses and synergy across the 4 × 4 Nav1.7/Nav1.8 subtraction matrix were generated for each cluster using the original rheobase values. All analyses were performed in Python using scikit-learn, NumPy, pandas, matplotlib, and seaborn.

### Statistics

Data are presented as mean ± SEM unless otherwise noted. Statistical comparisons were performed using appropriate parametric or nonparametric tests depending on distribution and experimental design, with significance set at p < 0.05. See Supplementary Table 1 for a description and results of all statistical tests. Cells that did not meet predefined electrophysiological quality criteria (unstable RMP and rheobase, overshoot less than +34 mV, in control) were excluded from analysis.

