## Supplementary Materials for "Combinatorial logic of Nav channels in nociceptor excitability: Different degrees of synergy define distinct neuronal groups"

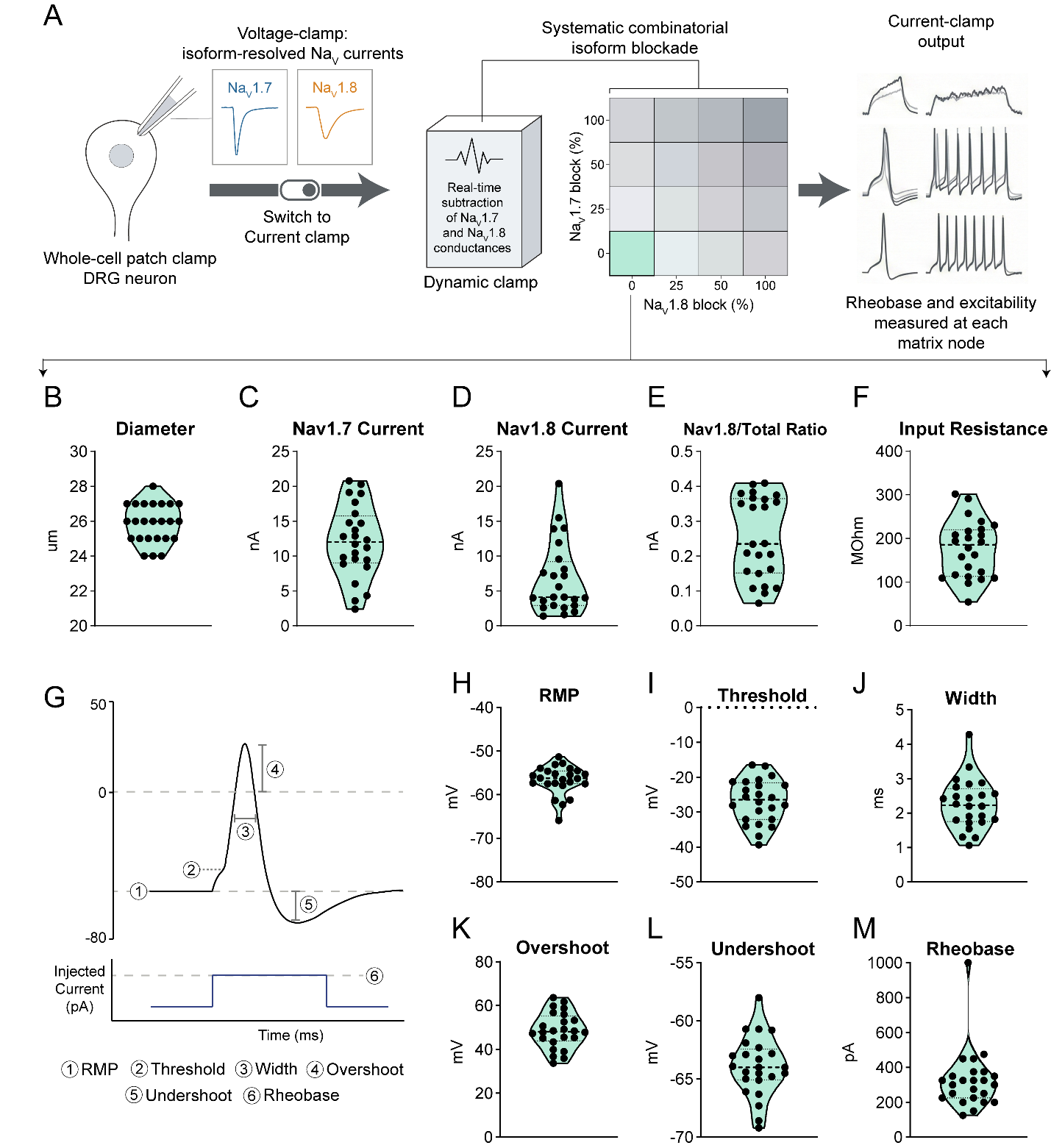
Supplementary Figure 1. Population-Wide Characterization of Baseline Biophysical Parameters

(**A**) Overview of the transitions from voltage-clamp characterization to the 16-node current-clamp matrix.

(**B**) Distribution of cell sizes within the sampled population, showing a focus on small-to-medium diameter neurons

(**C-E**) Baseline distribution for Nav1.7 and Nav1.8 currents as well as the Nav1.8/Total current ratio

(**F**) Input resistance distribution

(**G**) Action potential parameters measured included Resting Membrane Potential (RMP), Threshold, Width, Overshoot, Undershoot, and Rheobase. Population-wide distributions for RMP (**H**), Threshold (**I**), Action Potential Width (**J**), Overshoot (**K**), Undershoot (**L**), and Absolute Rheobase (**M**). These metrics establish the baseline state of the cells prior to simulated isoform blockade.

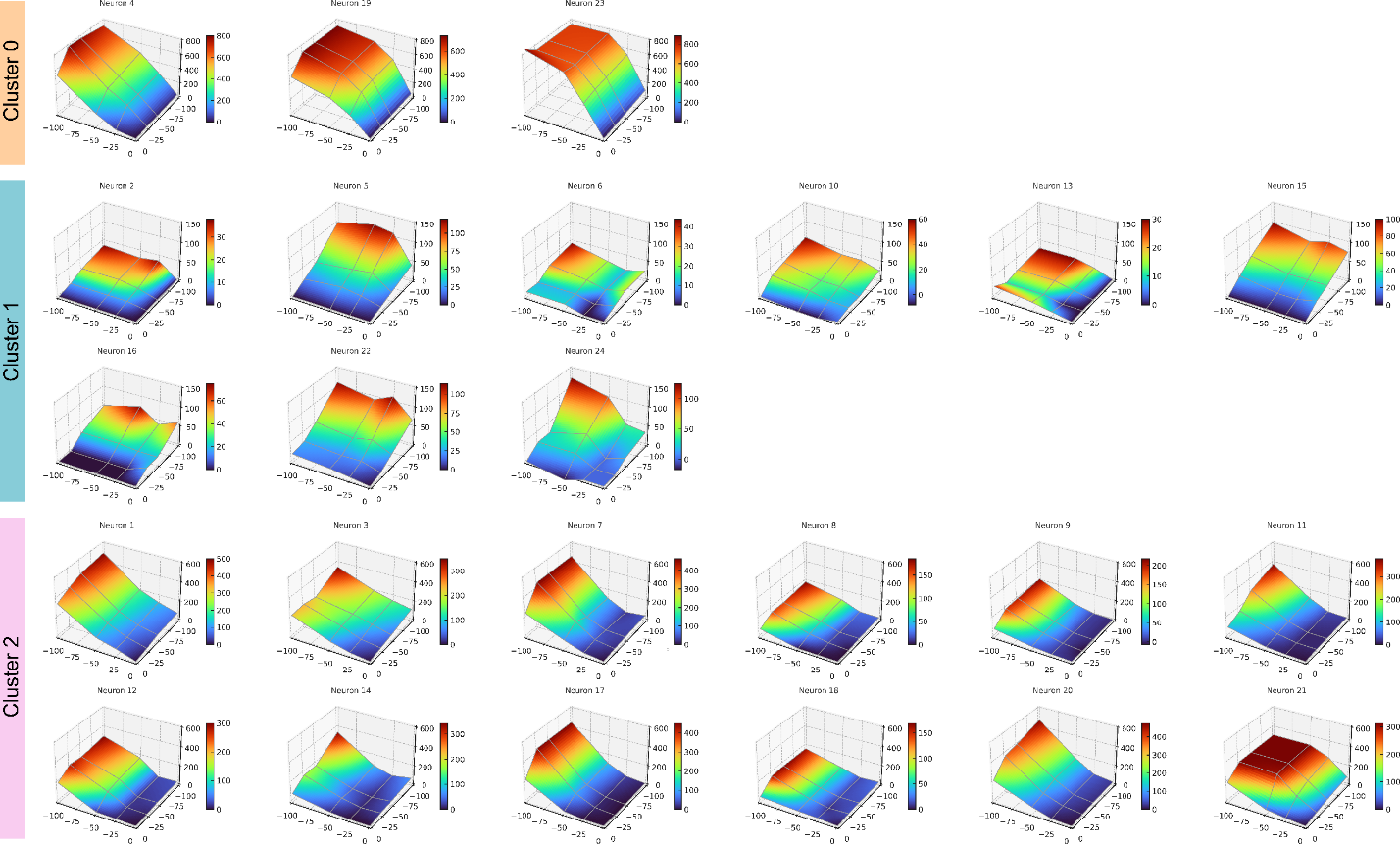

### Supplementary Figure 2. Individual cell heatmaps of rheobase response vectors

3D surfaces mapping increases in rheobase (rheobase change from control (%): 100*(Rheobase – Rheobase_Control)/Rheobase_Control %) across varying percentages of Nav1.7 and Nav1.8 block. Note the difference in surface directionality between clusters (labeled on the left).

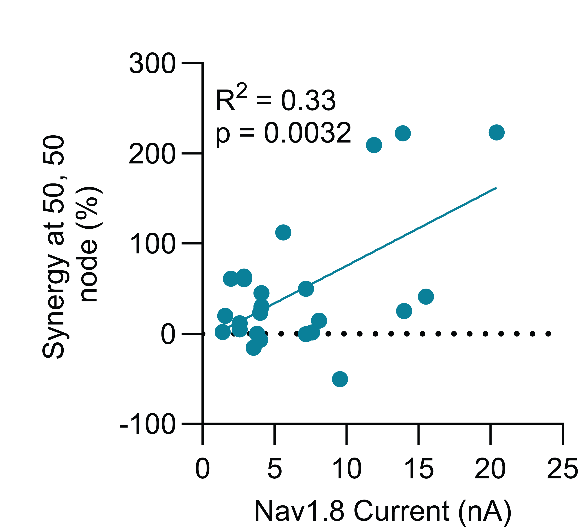

### Supplementary Figure 3. Linear regression of Nav1.8 current at baseline and synergy at the 50, 50 node

Nav1.8 current at baseline significantly correlates with the synergy value at the 50, 50 node but only explains 33% of the variation.

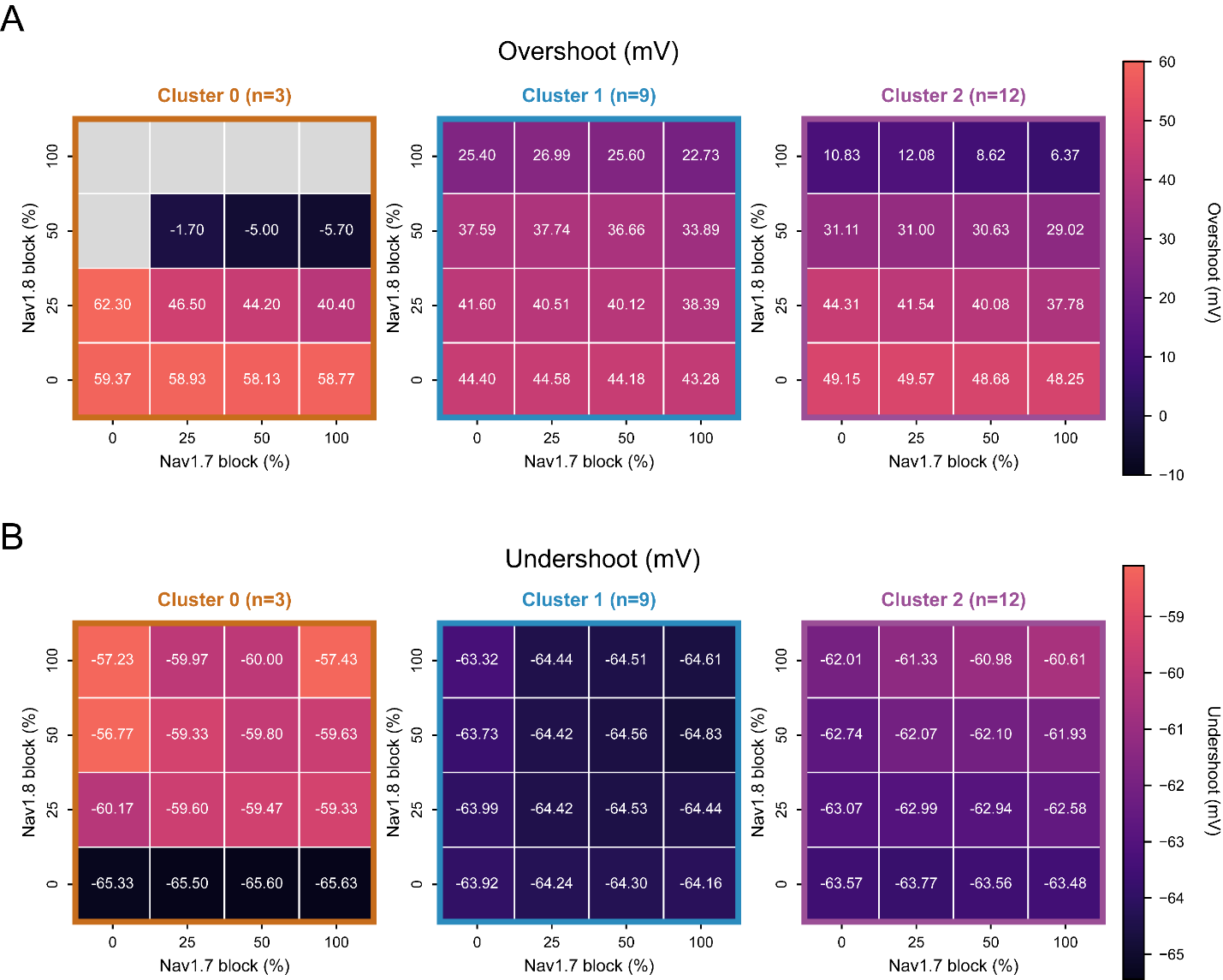

Supplementary Figure 4. Cluster-specific changes in action potential undershoot and overshoot across the Nav1.7 x Nav1.8 block matrix.

(**A**) Mean action potential overshoot and (**B**) mean action potential undershoot are shown across all combinations of Nav1.7 and Nav1.8 block for each cluster. Values represent cluster means at each matrix node. Cluster 0 cells showed the greatest reduction in overshoot and the largest shift in undershoot with increasing NaV block, whereas Cluster 1 cells were comparatively resistant. Cluster 2 showed an intermediate phenotype.

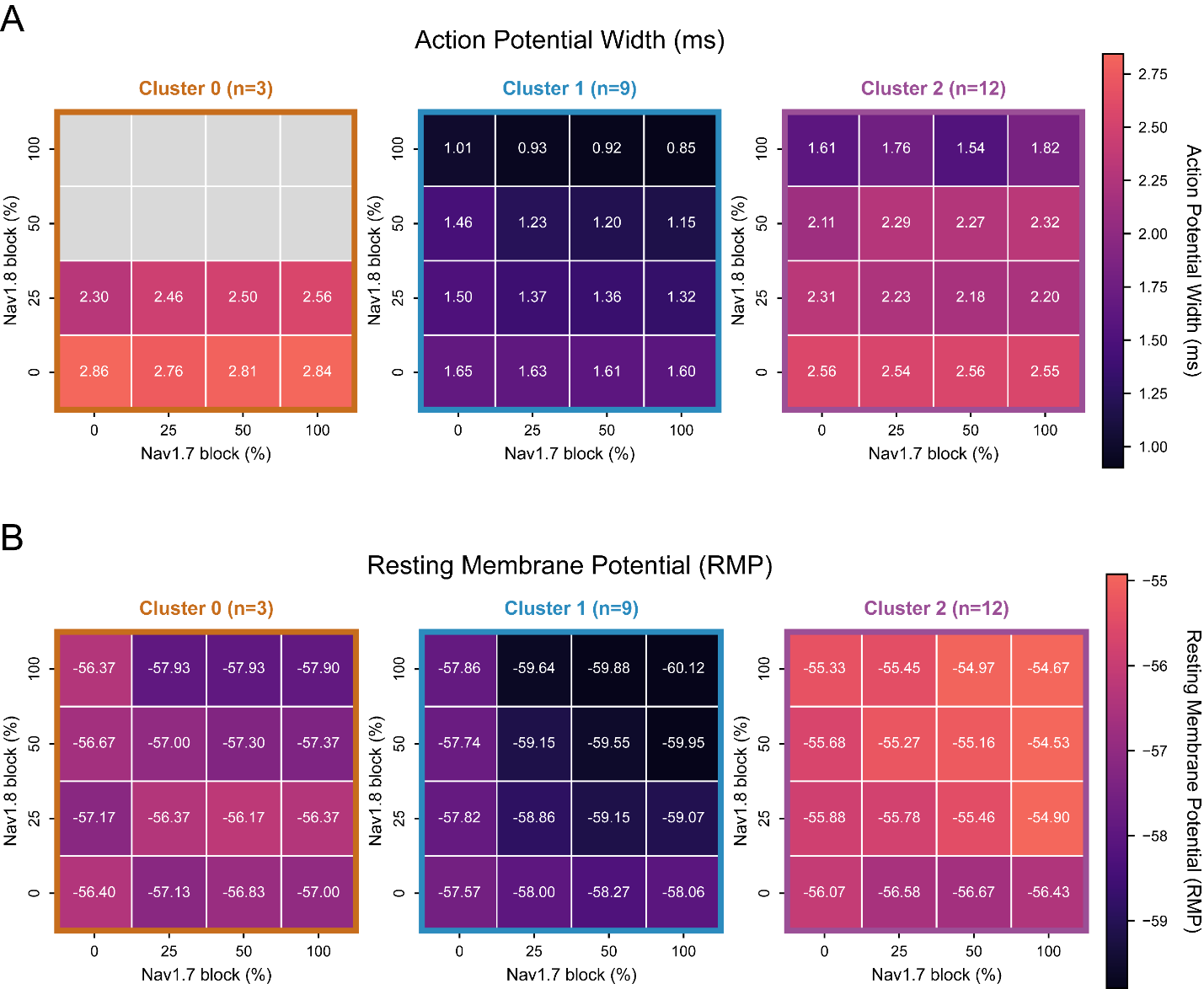

Supplementary Figure 5. Cluster-specific changes in action potential width and resting membrane potential across the Nav1.7 x Nav1.8 block matrix.

(**A**) Mean action potential width and (**B**) mean resting membrane potential (RMP) are shown across all combinations of Nav1.7 and Nav1.8 block for each cluster. Values represent cluster means at each matrix node.

### Supplementary Table 1. Statistical summary

| **Figure** | **Test type** | **Effect or comparison** | **p^a^** |
| --- | --- | --- | --- |
| Figure 2C | *Kruskal-Wallis test* |  | 0.1431 |
| Figure 2D | *Kruskal-Wallis test* |  | 0.0296 |
|  | *Dunn’s multiple comparison test* | |  |
|  |  | C0 vs C1 | 0.0274 |
|  |  | C0 vs C2 | 0.2624 |
|  |  | C1 vs C2 | 0.4493 |
| Figure 2E | *Kruskal-Wallis test* |  | 0.0027 |
|  | *Dunn’s multiple comparison test* | |  |
|  |  | C0 vs C1 | 0.0140 |
|  |  | C0 vs C2 | >0.9999 |
|  |  | C1 vs C2 | 0.0117 |
| Figure 2F | *Kruskal-Wallis test* |  | 0.0654 |
| Figure 2G | *Kruskal-Wallis test* |  | 0.0027 |
|  | *Dunn’s multiple comparison test* | |  |
|  |  | C0 vs C1 | 0.0201 |
|  |  | C0 vs C2 | >0.9999 |
|  |  | C1 vs C2 | 0.0158 |
| Figure 2G | *Kruskal-Wallis test* |  | 0.0027 |
|  | *Dunn’s multiple comparison test* | |  |
|  |  | C0 vs C1 | 0.0201 |
|  |  | C0 vs C2 | >0.9999 |
|  |  | C1 vs C2 | 0.0158 |
| Figure 2H | *Kruskal-Wallis test* |  | 0.0020 |
|  | *Dunn’s multiple comparison test* | |  |
|  |  | C0 vs C1 | 0.0151 |
|  |  | C0 vs C2 | >0.9999 |
|  |  | C1 vs C2 | 0.0071 |
| Figure 2I | *Kruskal-Wallis test* |  | 0.0350 |
|  | *Dunn’s multiple comparison test* | |  |
|  |  | C0 vs C1 | 0.0295 |
|  |  | C0 vs C2 | 0.1837 |
|  |  | C1 vs C2 | 0.7347 |
| Figure 2L | *Fisher’s exact test* |  | 0.4065 |
| Figure 4E | *Kruskal-Wallis test* |  | <0.0001 |
|  | *Dunn’s multiple comparison test* | |  |
|  |  | C0 vs C1 | 0.0004 |
|  |  | C0 vs C2 | 0.3010 |
|  |  | C1 vs C2 | 0.0023 |
| Supplementary Figure 3 | *Linear regression* |  | 0.0032 |

^a^adjusted p-values for multiple comparisons testing
